# A Universal Method for Crossing Molecular and Atlas Modalities using Simplex-Based Image Varifolds and Quadratic Programming

**DOI:** 10.1101/2023.03.28.534622

**Authors:** Kaitlin M. Stouffer, Alain Trouvé, Laurent Younes, Michael Kunst, Lydia Ng, Hongkui Zeng, Manjari Anant, Jean Fan, Yongsoo Kim, Michael I. Miller

## Abstract

This paper explicates a solution to the problem of building correspondences between molecular-scale transcriptomics and tissue-scale atlases. The central model represents spatial transcriptomics as generalized functions encoding molecular position and high-dimensional transcriptomic-based (gene, cell type) identity. We map onto low-dimensional atlas ontologies by modeling each atlas compartment as a homogeneous random field with unknown transcriptomic feature distribution. The algorithm presented solves simultaneously for the minimizing geodesic diffeomorphism of coordinates and latent atlas transcriptomic feature fractions by alternating LDDMM optimization for coordinate transformations and quadratic programming for the latent transcriptomic variables. We demonstrate the universality of the algorithm in mapping tissue atlases to gene-based and cell-based MERFISH datasets as well as to other tissue scale atlases. The joint estimation of diffeomorphisms and latent feature distributions allows integration of diverse molecular and cellular datasets into a single coordinate system and creates an avenue of comparison amongst atlas ontologies for continued future development.

## 1 Introduction

Since the 17th century, scientists have seen living organisms as a hierarchy of biological mechanisms at work across scales. To understand the interplay of these mechanisms, reference atlases that incorporate genetic, cellular, and connectivity measures into a single coordinate space have been constructed and which aim to summarize the mass of data across scales through a set of discrete partitions. An instance of the more general segmentation problem in computer vision, atlas construction relies on the underlying assumption of homogeneity within each region. The optimal partitioning assigns a label to each region based on this homogeneity and the presence of sharp changes at the boundaries between regions.

In biology, this label frequently reflects behavior or function, as seen in two of the most common mouse brain atlases: the Allen Reference Atlas (ARA) [1] and the Franklin and Paxinos Atlas [2]. Together, the common coordinate framework an atlas provides in addition to its ontology have guided research efforts in facilitating the comparison of different types or replicates of data in a single coordinate system and in honing efforts of study to particular regions relevant to each unique investigation.

The widespread use of these atlases, particularly in the fields of digital pathology and neuroimaging, has motivated efforts to develop image registration tools to align individual images to such reference atlases. A large family of methods, all diffeomorphism based [3], have been developed within the field of Computational Anatomy (CA) [4, 5] for transforming coordinate systems at the tissue scales. These come particularly from multiple labs in the magnetic resonance imaging (MRI) community [6, 7, 8, 9, 10, 11, 12, 13, 14]. More recently, Molecular CA [15] has emerged which unifies the dense tissue scales of MRI with high resolution micron scales of digital pathology imagery. These approaches are hierarchical [16], constructing what are termed image varifold representations, which are geometric measures of the brain existing at multiple scales, and therefore allowing for the simultaneous representation of both micron scale particle phenomena, such as the transcriptional or cell type data studied herein, and millimeter scale tissue phenomena, as traditionally studied in CA.

A central challenge that remains within these representations is the necessity of crossing between those reflecting different imaging modalities and therefore different functional range spaces, which exist at different scales. In the setting of classical images on a regular grid, this challenge has been addressed through different approaches including matching based on analytical methods using cross-correlation [17] or localized texture features [18], and methods for transforming one range space to another in crossing modalities and scales based on polynomial transformations [19], scattering transforms [20] and machine learning [21, 22, 23]. More recently, methods in deep learning have also been applied to align single-cell datasets, modeled as regular grid images, both to atlases at the histological scale [24] as well as reference transcriptional atlases that are beginning to emerge [25].

We should however expect even more diversity in the types and scales of data that can be measured with the rapidly developing technologies in imaging and spatial transcriptomics. These aim to detect up to thousands of genes simultaneously with spatial information, and thus, allow us to view both the micro and even nanometer scales with exquisite detail [26]. Both the diversity and magnitude of this data pushes the limits of our ability to model such datasets as classical continuous images, discretized on regular grids. Indeed, as seen in those repositories generated in the BRAIN Initiative Cell Census Network (BICCN) and archived at the Brain Image Library (BIL), these datasets are already on the order of terabytes and will only continue to increase as technologies shift from mouse to human measurements. Hence, the need remains for a modeling framework equally equipped to represent datasets in both forms of traditional continuous imagery sampled on a regular grid, and those of discrete particles with attached functional description; and for an associated registration mechanism to align objects in this framework across different functional modalities at different scales.

This paper focuses on the use of mesh-based image varifolds, as described in [27], for simultaneously modeling molecular and tissue scale data. A subproblem covered by the mesh based image-varifold theory outlined in [27] is the mapping of molecular scale data to atlas coordinate systems. Image varifolds are geometric measures [15], which allow us to provide a single representation that supports molecular transcriptomics measurements, cell-based measurements, and tissue scale atlases. We explicate, here, the construction of a universal method rooted in this framework for transferring molecular scale data to tissue scale “cartoon” atlases, which are devoid of gene measurements, and rather, only contain a fixed partition into structures in their description.

Our solution couples coordinate system transformation via geodesic generation of minimal energy diffeomorphisms to estimation of a family of probability laws, which give for each atlas label, a distribution over molecular features that is the most reasonable explanation of the target transcriptomic dataset. Specifically, we model each atlas region as homogeneous and stationary with respect to space, giving an optimal alignment between atlas and target that maximizes similarity in distribution over features across each site in a single atlas region while minimizing the energy of the geometric deformation (diffeomorphism). This consequently skews emphasis away from the foreground-background boundaries that almost exclusively govern image alignment and instead highlights the underlying assumptions in the architecture of the cartoon atlas, whose boundaries were initially constructed so as to maximize the homogeneity of the region. We estimate the diffeomorphism and probability laws jointly via an alternating algorithm, as explicated here, that iterates large deformation diffeomorphic metric mapping (LDDMM) with quadratic programming for minimizing the normed distance between the template and target, and as a result, yields both spatial alignment and functional correspondence between template and target.

We demonstrate the efficacy of this methodology in mapping 2D sections of the ARA [1] to corresponding sections of both cell-independent and cell-based spatial transcriptomics datasets, both generated via the MERFISH imaging-based spatial transcriptomics technology, which yields single molecule resolution. We present methods for sparsifying the functional transcriptome descriptions via gene selection based on mutual information with spatially discriminating variables and subsequently illustrate the stability of our estimated diffeomorphisms to choices of subsets of features. Finally, given the plethora of existing reference atlases, each of which might define a different partitioning scheme over the same area of tissue, questions of comparison and relevance of each atlas to emerging molecular and cellular signatures naturally arise [28]. We show through the use of our methodology to map not just atlas to molecular dataset, but one atlas to another, that the correspondence yielded by our method serves as an anchor for re-examining existing ontologies and creating new ones for the future.

## 2 Results

### 2.1 Image Varifolds and Transformations for Molecular Scales Based on Varifold Norms

In Computational Anatomy, correspondence between tissue sections is computed using coordinate transformation between the sections by solving an optimization problem characterized by the set of possible transformations to optimize the image similarity function that specifies the alignment of the sections. These transformations are modeled as affine motions and diffeomorphisms *φ* which act to generate the space of all configurations. For classical images such as for MRI, LDDMM [29] uses the action of diffeomorphisms on images *I* as classical functions using function composition on the right with the inverse of the diffeomorphism: *φ* · *I*(*x*) = *I* ○ *φ*^-1^(*x*) for *x* ∈ *R^d^*. The image similarity function used is often a norm on functions, and solving the problem of minimization of the norm in the space of diffeomorphisms gives the metric theory of LDDMM for generating geodesic matching between exemplar anatomies [30, 31].

Spatial transcriptomics generates measurements that while often represented as regular lattice images, are fundamentally lists of point measurements across the different technologies and thus, often dispersed irregularly over space. In spot-resolution technologies including Visium, DBiTseq, and SlideSeq, these point measurements are the magnitudes of gene expression in the neighborhood of each “spot”, which could be placed in a regular grid pattern. In contrast, in imaging-based spatial transcriptomics technologies including STARmap, Barseq, SeqFISH, and MERFISH, as illustrated here, these point measurements are single mRNA molecules or single cells, therefore dispersed in space according to the given tissue architecture and instantaneous cell dynamics measured. In both cases, we can represent these point measurements as “particles”.

Natural fluctuation in gene expression over time and space coupled to the dynamics of each spatial transcriptomics technology leads each tissue section, at the molecular (1-100 micron) scale, to have a varying number of such particles with no natural ordering of particles consistently apparent between sections. To build correspondences between these datasets of point measures, we unify the molecular scales with image-like functions as has been developed for building correspondences at tissue scales in MRI [4]. For this we represent the particles as “generalized functions” [15]. Since they carry gene or cell image data we call them image varifolds [27], linking to the rich literature on the geometric measure theory of varifolds. This allows us to represent particle clouds at any scale in both spatial and imaging function dimensions. We note landmark-based methods [32] that assume direct permutation correspondence between particles across images are not applicable, as a MERFISH section may have 100,000 particles requiring an unfathomable number of permutations to specify.

Varifolds are defined as follows. We consider a Euclidean space in d dimensions with *d* = 2, 3, to which we add function dimensions represented by a set 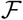. In spatial transcriptomics datasets, the functional dimensions represent the gene types of detected mRNA transcripts, treated as independent measures or aggregated into cells or small neighborhoods. At the finest scale, we model a discrete set of point measures (particles) reflecting the individual reads recorded by the given technology, whether they be single transcripts or distributions of transcripts in a given cell or neighborhood. To a single read 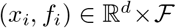, we associate the elementary “Dirac” measure, *δ_x_i__* ⊗ *δ_f_i__*, which acts on a set 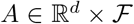 as *δ_x_i__* ⊗ *δ_f_i__* (*A*) = 1 if (*x__i__, f_i_*) ∈ *A* and 0 otherwise. The point measures carry weights *w_i_*, giving the multiplicity, typically as number of transcripts or number of cells measured by each individual read. The discrete image varifold is defined as the weighted sum of Diracs representative of the collection of particles and functional features (*w_i_, x_i_, f_i_*), *i* =1, 2 …:

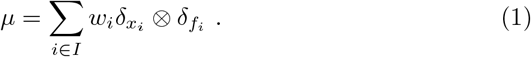

While for the molecular scales, each data point is a measurement of a single mRNA transcript or local (e.g. cell’s) distribution on the feature space of gene type 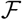, in contrast, a data point in a given atlas is interpreted as a single voxel with a label prescribed to it from the overall ontology, 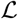.

It is natural to associate a density in mass per unit volume to the varifold through the classical decomposition of measures as a product. This gives the marginal distribution *ρ* on physical space, 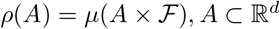, *A* ⊂ ℝ^*d*^, and the field of conditional probability measures over the feature space *μ_x_, x* ∈ ℝ^*d*^ on 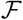 the feature space:

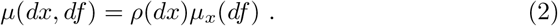

For molecular scales, *ρ*(·) on ℝ^*d*^ is typically the spatial distribution of total gene expression, while for atlas images at tissue scales, it is a continuous uniform distribution over the support of the tissue. Cross-modality mappings from molecular to tissue scales thus imbue the atlases with estimates *ρ*(·) and the field of conditional probabilities *μ_x_*(·), *x* ∈ ℝ^*d*^ of the molecular feature space (e.g. gene type).

### 2.2 Quadratic Program for Cross Modality Mapping on Meshes

A central goal is to imbue the atlas with molecular or cellular information by estimating a cross-modality mapping between the atlas and a finer scale, single-cell or subcellular dataset, such as those emerging particularly from imaging-based spatial transcriptomics technologies. To compute this mapping, we model each modality as an image varifold, a product of measures over physical and feature space, by instantiating each measurement as a triangulated or simplex mesh following [27]. Each mesh carries a collection of vertices **x** = (*x_i_* ∈ ℝ^2^)_*i*∈*I*_. From the vertices we construct the simplex triangles *γ_j_* (**x**) and their centroids *m_j_*(**x**) for *j* ∈ *J*, with vertex numbers |*I*| and simplices | *J*| determined by the resolution selected.

We denote the target mesh as *τ* throughout the paper; see Section 4.1 for detailed construction. We note the triangles and centers are a function of the underlying vertices, but we will often suppress their explicit dependence except when necessary. To complete the image varifold we append to the mesh *τ* the density ***α*** = (*α_j_*)_*j*∈*J*_ and the field of probability laws ***ζ*** = (*ζ_j_*)_*j*∈*J*_ on 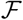:

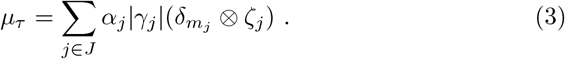

Importantly, in spite of the apparent differences between equations (1), (2) and (3), they all belong to the same category of mathematical objects, and can be addressed together in the framework of image varifolds.

At the molecular scales presented in this paper, the density is number of cells or number of mRNA transcripts per mm^2^ and is defined for each simplex, area *I*, as

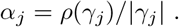

The field *ζ_j_, j* ∈ *J* are probabilities over genes or cell types with finite dimensional feature spaces 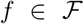, with 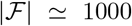 in the case of genes and 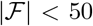 for cell types. Each *ζ_j_* (*f*) is a probability of gene or cell type, with 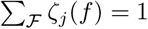, indexed by location in the image.

We take the ARA [1] as the template to be mapped onto the molecular data. For mapping the atlas to molecular scales, we have to estimate both the diffeomorphisms *φ* : ℝ^*d*^ → ℝ^*d*^ transforming atlas coordinates. as well as the unknown densities and conditional feature distributions, ***α***^*π*^, ***ζ***^*π*^ which we take as latent variables for the atlas. We denote the mesh for the template as *τ*_0_ representing its vertices **x**^0^ = (*x_i_*)_*i*∈*I*_0__ and simplices and centers (*γ_j_, m_j_*), *j* ∈ *J*_0_.

The atlas carries a finite ontology, 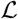, dividing it into disjoint spatial partitions. We model each atlas region as having a distribution (non-normalized) over the molecular features 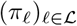 on 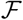 viewed as latent variables that are homogeneous across the partition region. The simplex law is determined by the contribution of each ontology region to the vertex for *j* ∈ *J*_0_, given by the mixture distribution 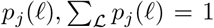. The atlas has appended the molecular feature space estimated from the target (***α***^*π*^, ***ζ***^*π*^) and is given as:

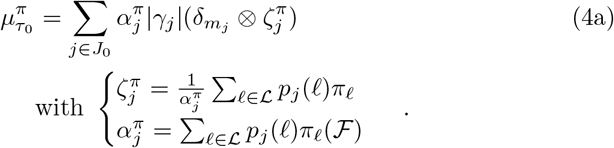

The group action carrying the atlas onto the target becomes

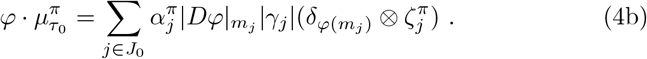

Here, |*Dφ*|_*m_j_*_ is the Jacobian determinant of *φ* at *m_j_*.

Figure 1 shows a mesh-based image varifold for two coronal sections (*Z* = 385, *Z* = 485) of the Allen Atlas, with finest granularity ontology 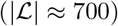 and with meshes rendered at 50 *μ*m resolution. The right panel of Figure 1 illustrates the physical densities *α_j_* ∈ *J* of mRNA transcripts per mm^2^ in coronal sections of MERFISH from the Allen Institute [33].

**Fig. 1.**
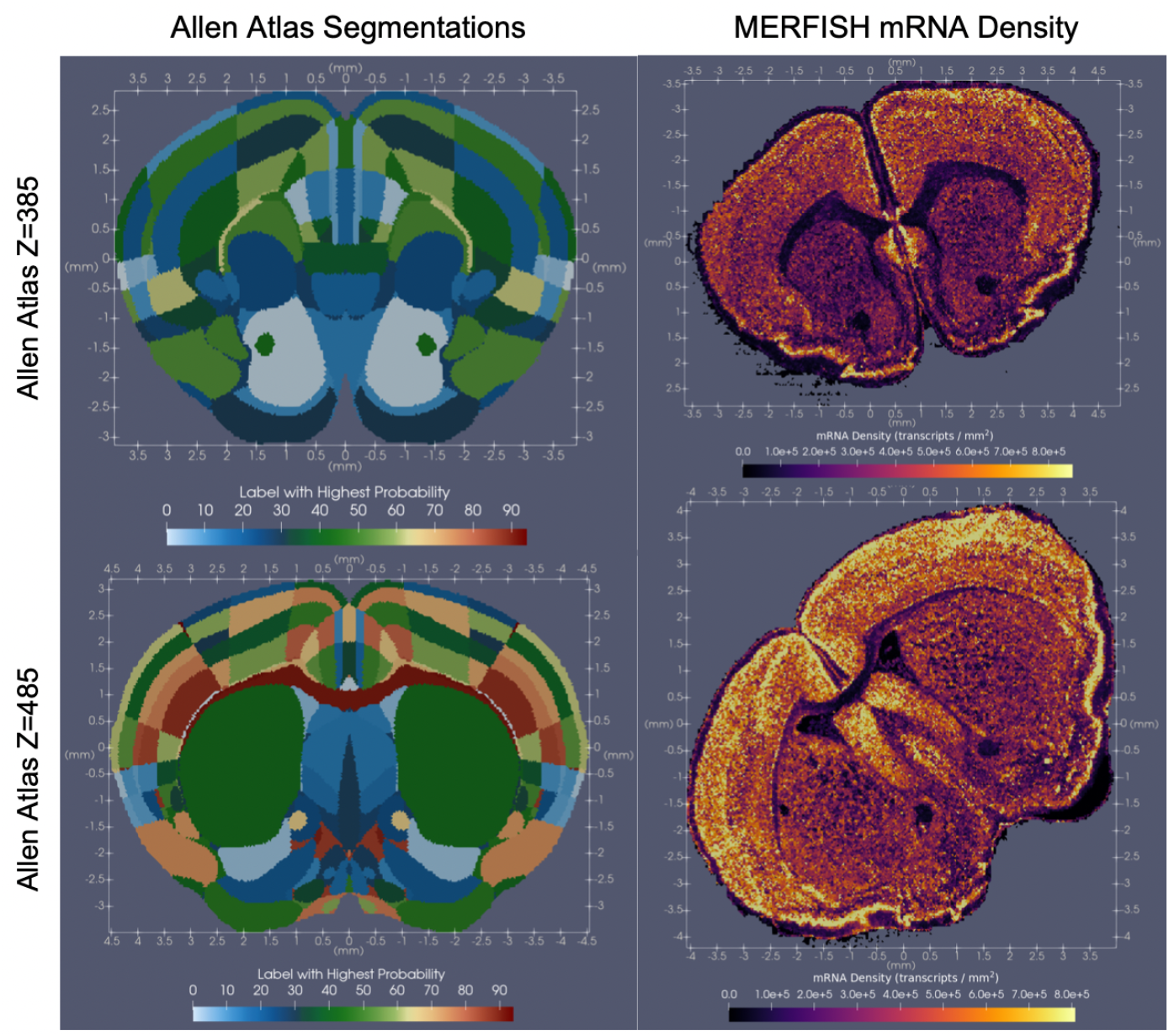
Coronal sections of mouse brain rendered as mesh from Allen Reference Atlas (left) and MERFISH-spatial transcriptomics (right). Selected sections of atlas chosen by visual inspection to match MERFISH architecture. Meshes are rendered at 50 *μ*m, with tissue sections corresponding to Z-sections 385 (top row) and 485 (bottom row) in 10 *μ*m Allen reference atlas. Colors in the left column indicate a region in the Allen ontology, while colors in the right column indicate the density of mRNA transcripts given by the number of transcripts per simplex area *α_j_* ≠ # transcripts/|*γ_j_*|.

To map the mRNA measures to atlases we follow [27] and define the space of image varifolds *μ* ∈ *W*\* to have a norm 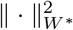, and transform the atlas coordinates onto the targets to minimize the norm. The space of varifold norms is associated to a reproducing kernel Hilbert space [34, 15] (see (7) below) defined by the inner-product of the space as 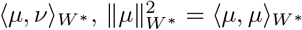.

The mapping variational problem constructs *φ* : ℝ^*d*^ → ℝ^*d*^ and feature laws 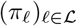 on 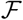 to carry 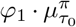 onto *μ_τ_* minimizing the normed difference. Densities that are estimated are constrained to fall in the range 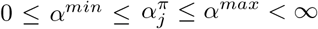 to ensure positive values for the density and incorporate prior knowledge of cellular or molecular distributions.

#### Variational Problem 1

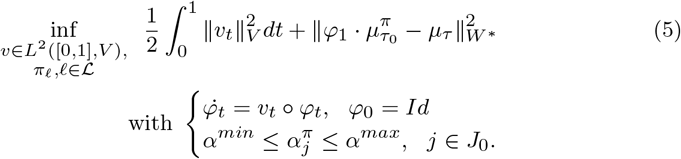

Throughout, we take *α^min^* to be the 5th percentile of values of (*α_j_*)_*j*∈*J*_ in the target.

The variational problem maximizes the overlap of homogeneous regions in the atlas (e.g. each partition in the ontology) with those in the target (e.g. regions where conditional feature distributions are stationary over space) by deforming coordinates using LDDMM to optimize the vector field *v_t_*, *t* ∈ [0,1]. The quadratic programming calculations solve for *π_i_*, 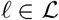 for the atlas to gene expression and cell-type problem and are described in the methods section 4.3.

Figure 2 illustrates the results of mapping the Allen Atlas coronal sections to Allen MERFISH spatial transcriptomics sections, shown in Figure 1. Allen Atlas sections were chosen based on correspondence through visual inspection. Estimated mRNA densities, 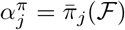, as depicted on left and middle columns, were achieved through solution of the quadratic program as defined in (9), and reflect total mRNA densities from a full set of 702 genes as features. Leftmost column shows estimated mRNA densities on transformed geometry of atlas mesh under the action of the diffeomorphism *φ*, while middle column shows estimated mRNA densities on original atlas geometry. Right column shows the action of the diffeomorphism *φ* on each atlas section, with vertex positions, *φ*(***x***^0^), and with approximate determinant jacobian, |*Dφ*|*_m_j__* indicated by the color at each simplex site *j* ∈ *J*_0_.

**Fig. 2.**
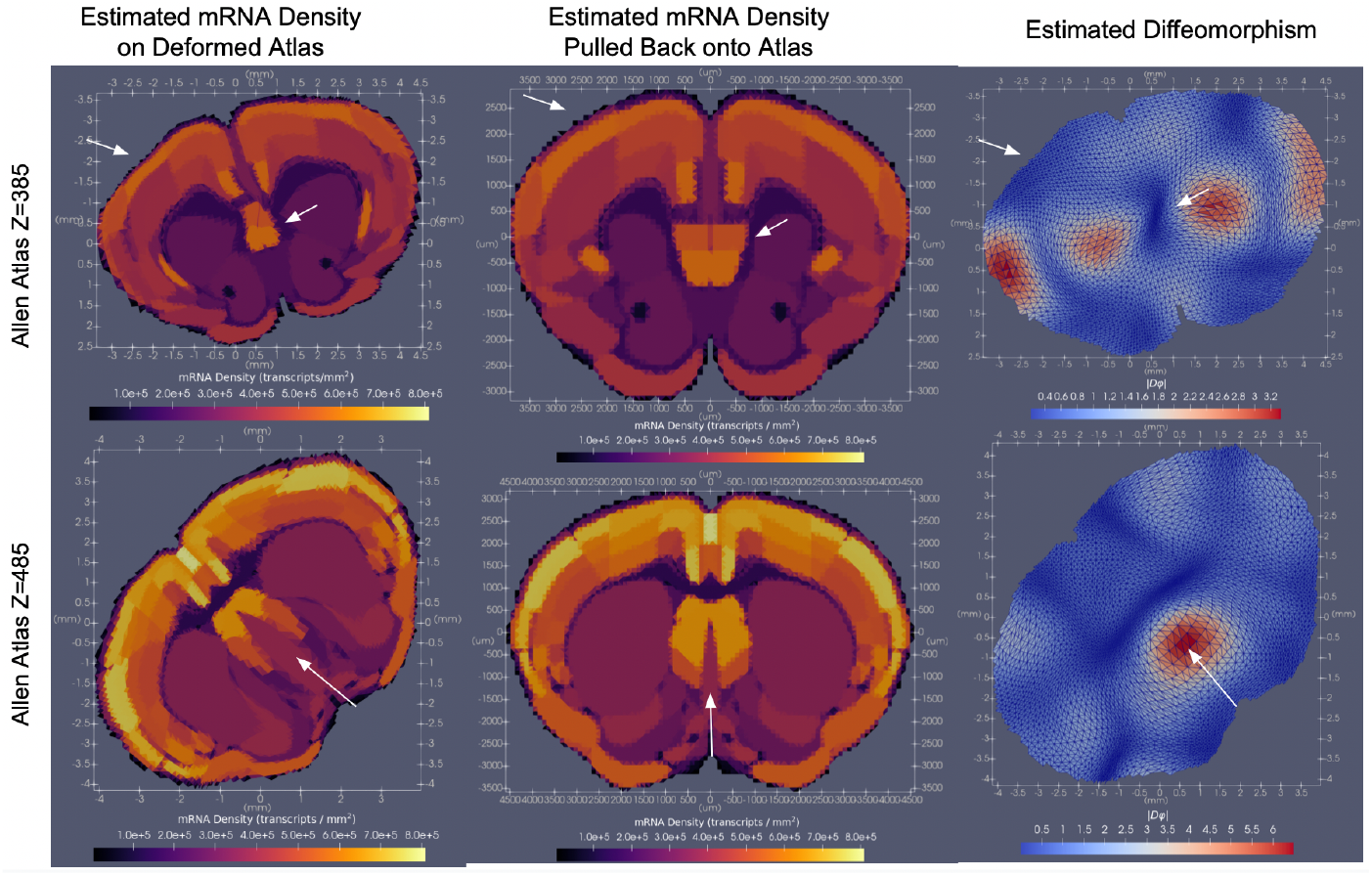
Results of cross-modality atlas mapping to Allen MERFISH spatial transcriptomics [33] for coronal sections of tissue at approximate Allen atlas Z-sections of 385 (top) and 485 (bottom). Left column shows estimated mRNA densities, 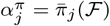, *j* ∈ *J*_0_, per deformed simplex site under the action of the diffeomorphism of atlas to target space 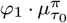; middle column shows the same pulled back onto original atlas geometry 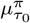; right column shows the diffeomorphism applied to the mesh *τ*_0_, with depicted approximation of the determinant of the Jacobian |*Dφ*_1_|*_m_j__*, *j* ∈ *J*_0_, as described in Section 4.3.

### 2.3 Dimension Reduction of Gene Distributions via Mutual Information

In mapping atlases to distributions of mRNA, we are typically interested not just in overall mRNA density, but the distribution of expression across a particular set of genes. The size of the total gene set measured varies across technologies, ranging from hundreds to tens of thousands of different genes [26]. However, both computational time and memory frequently dictate the analysis of only a subset of these genes at a time, together with their relevance to each particular application. A common selection mechanism is to consider those genes that are most “spatially variable” [35] or “differentially expressed” [36], under the assumption that expression pattern thereby varies per biologically different regions of tissue. This is particularly relevant, here, in the context of mapping spatial transcriptomics to atlases where we aim to estimate distributions over genes for each region in our atlas that we assume is homogenous within the region.

Various methods have been described for identifying which genes in a spatial transcriptomics dataset are more spatially varying than others, some examples being Gaussian process registration, Laplacian Score, [35] and Moran’s I [37]. In order to score genes which are most spatially varying we introduce Mutual Information scoring which assesses the differential expression of genes in space in a cell-independent manner. Specifically, we score each gene with the mutual information between the two random variables *X, M* which capture, respectively, an orientation in space and a relative density of mRNA expression for that gene (see Section 4.4). In the case of serial sections, as in the MERFISH data from the Allen Institute, each gene is assigned a score per section, with tallies taken across all sections to deduce which genes are most spatially variable across the entire brain. We note this approach is similar in spirit but not identical to that in [36] which uses the Kullback-Leibler divergence to find genes with differential expression across cells distributed in space.

Shown in Figure 3 is a single section of Allen Institute MERFISH data depicting the distribution of three example genes with the lowest mutual information scores (top row, *Chodl, Brs3, Hpse2*) and the highest mutual information scores (middle row, *Gfap, Trp53i11, Wipf3*) computed across the whole set of 60 serial sections. In each case, conditional probabilities, *ζ_j_* (·) reflect the relative occurrence of each gene in the context of a subset of 20 total genes of either lowest (top row) or highest (middle row) mutual information. In line with expectations, 75% of the genes comprising those with scores in the bottom 25% of the total 700 genes were decoy genes (e.g. ‘BLANK’) without biological meaning but used as controls for assuring the quality of the dataset.

**Fig. 3.**
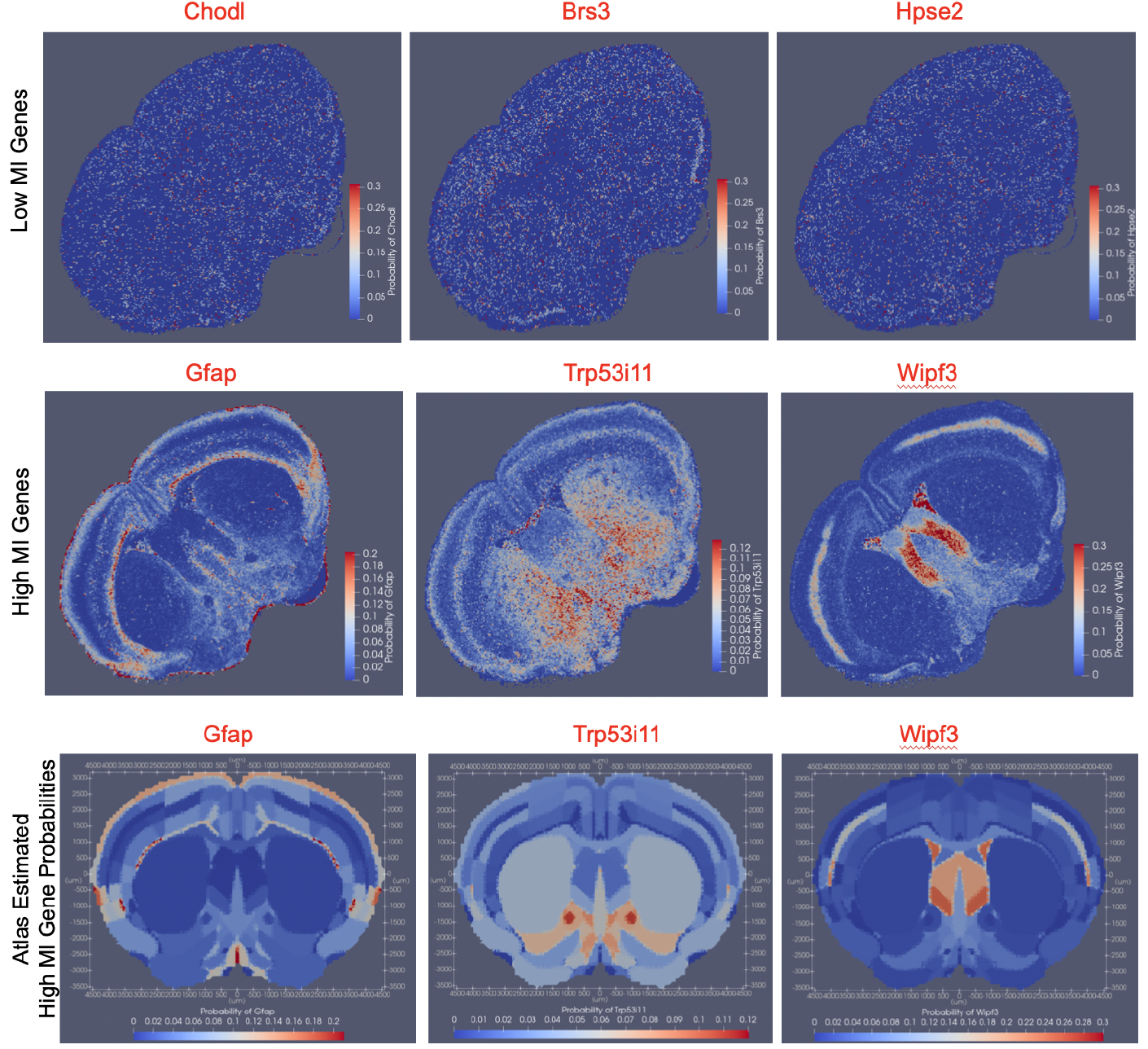
Relative expression per simplex, *ζ_j_* (·), of three genes with lowest (top row) and highest (middle row) mutual information score, computed across the entire set of 60 coronal sections in Allen Institute MERFISH sample, and shown on one section at approximately the coronal slice level of *Z* = 485 in the Allen atlas. Estimated probabilities 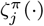 for each of the three genes with highest mutual information (*Gfap, Trp53i11, Wipf3*) shown for each atlas region with the native atlas geometry (bottom row).

Shown in the bottom row of Figure 3 is the estimated probability, 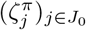, for each of the three genes with highest mutual information score (*Gfap, Trp53i11, Wipf3*), shown for each region on the Allen atlas section. For calculating these estimates we solve the variational algorithm with LDDMM and quadratic programming estimation of the gene feature distributions to map the Allen atlas section, *Z* = 485 to the Allen MERFISH target image-varifold . For this result, the smaller feature space of 20 total gene types corresponding to those with the highest mutual information scores are used for the mapping algorithm.

### 2.4 Mapping of Cell Distributions to Atlases

The two previous sections presented results solving the mapping problem between atlas and MERFISH based on the mRNA reads directly. Alternatively, these raw mRNA reads can be segmented into discrete cells as a mode of data reduction followed by downstream analyses clustering the cells into discrete cell types. The mesh-based image varifold framework is ideal for taking the measure representation directly on the aggregated cells and solving the variational problem of mapping to atlas coordinates. Figure 4 shows the results of mapping an Allen atlas section at *Z* = 675 to a section of cell-segmented MERFISH transcriptional data (courtesy of the JEFworks Lab, Johns Hopkins University). The total gene set measured is ≈ 500 genes, with each transcript assigned to a single cell. Transcriptional profiles per cell are clustered into 33 distinct clusters using Leiden graph-based clustering [38] and annotated as cell types based on known marker genes. This gives a cell-based dataset analogous to the transcript-based dataset discussed in Section 2.2 in which densities, *α_j_* (·), reflect the spatial density of data points (here, 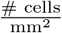), and conditional probability distributions, *ζ_j_*(·), are defined over the feature space of cell types, 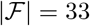.

**Fig. 4.**
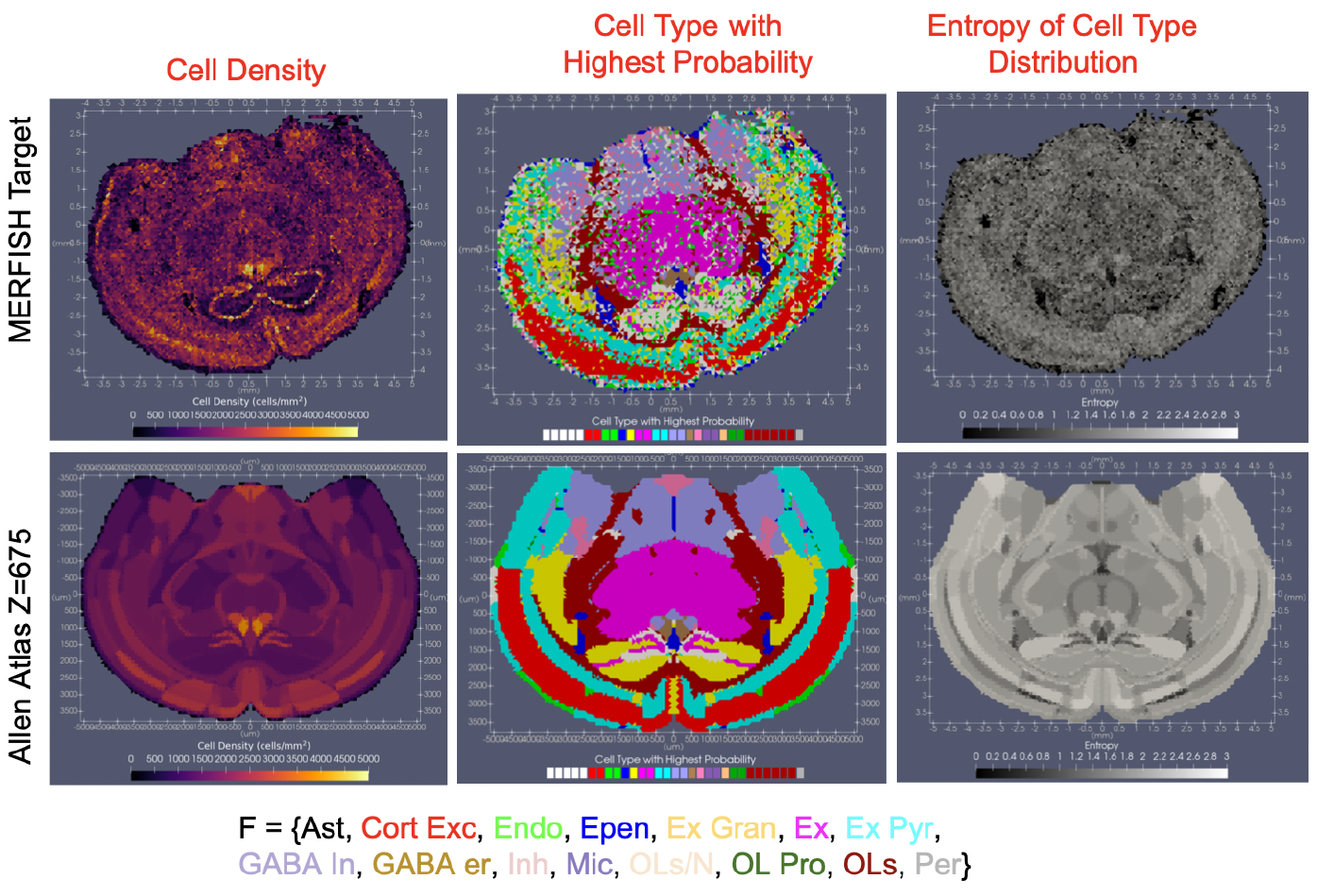
Cell densities and cell-type probabilities for MERFISH sections (top row). Estimated cell densities and cell type probabilities, 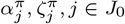, *j* ∈ *J*_0_ in Allen atlas section *Z* = 675 (bottom). Spatial density of cells given in units of 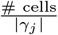 (left) and cell type probabilities summarized by depiction of cell type with highest probability for each simplex (middle), and entropy of probability distribution over cell types for each simplex (right). Specific subtypes of cell types (e.g. astrocytes type 1, astrocytes type 2, assigned same color according to labels shown in bottom of figure). Abbreviations of cell types: Astrocytes (Ast), Cortical Excitatory Neurons (Cort Exc), Endothelial Cells (Endo), Ependymal Cells (Epen), Excitatory Granule Cells (Ex Gran), Excitatory Neurons (Ex), Excitatory Pyramidal Neurons (Ex Pyr), GABAergic inhibitory neurons (GABA In), GABAergic Estrogen Receptor Neurons (GABA er), Inhibitory Neurons (Inh), Microglia (Mic), Oligodendrocytes / Neurons (OLs/N), Oligodendrocyte Progenitor Cells (OL Pro), Oligodendrocytes (OLs), Pericytes (Per).

The essential part of the model for estimating the atlas distribution over cell types is the stationarity of the model across each atlas partition. It is therefore natural to examine the entropy of the distribution within each atlas compartment as a measure of the multiplicity of cell types within a compartment. Shown in the right column is the entropy of the estimated probability distribution over cell types for each simplex in both target and atlas, with nonzero probabilities assigned to ~ 3 – 5 distinct cell types in each simplex of the target versus ~ 1 – 20 cell types in each simplex of the atlas, varying per region in the original ontology.

We emphasize that there are various methods for solving the segmentation to cells and thereby dimension reduction as determined by the specific imaging technology. Some of the methods are rooted in image-based segmentation schemes such as the Watershed algorithm, operating jointly on transcriptional data and immunofluorescence images such as DAPI stains [39], while others utilize learning-based methods [40] for accommodating often a wider diversity of cell shapes and sizes. In either case, the assignment of mRNA reads to specific cells introduces a layer of functional information at the micron scale, which can now be modeled in lieu of or in tandem with the functional information at the nanometer scale (e.g. raw mRNA reads) as the feature space of a target image varifold to which we wish to map sections of an atlas. The image-varifold method is universal in the sense that it is agnostic to which discrete object is forming the information that provides the substrate for building correspondence.

In addition to cell type as the features associated to the cell aggregated transcriptome data, the feature space can remain gene type generated by aggregating the individual mRNA transcripts into an average gene expression feature per cell across the span of tissue. Shown in Figure 5 are the distribution of two genes (*Ntrk3, Fzd3*) out of a subset of 7 chosen to have the highest mutual information score. By normalizing the total mRNA per cell to 1, we estimate for an atlas section, a density, ***α***^*π*^ in units of 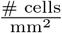, and a conditional distribution over gene types, ***ζ***^*π*^, reflecting the probability per cell in the given simplex, of mRNA belonging to each gene type. Figure 5 shows these estimated probabilities ***ζ***^*π*^ for those genes whose probability of expression per cell is correspondingly shown in the MERFISH target section.

**Fig. 5.**
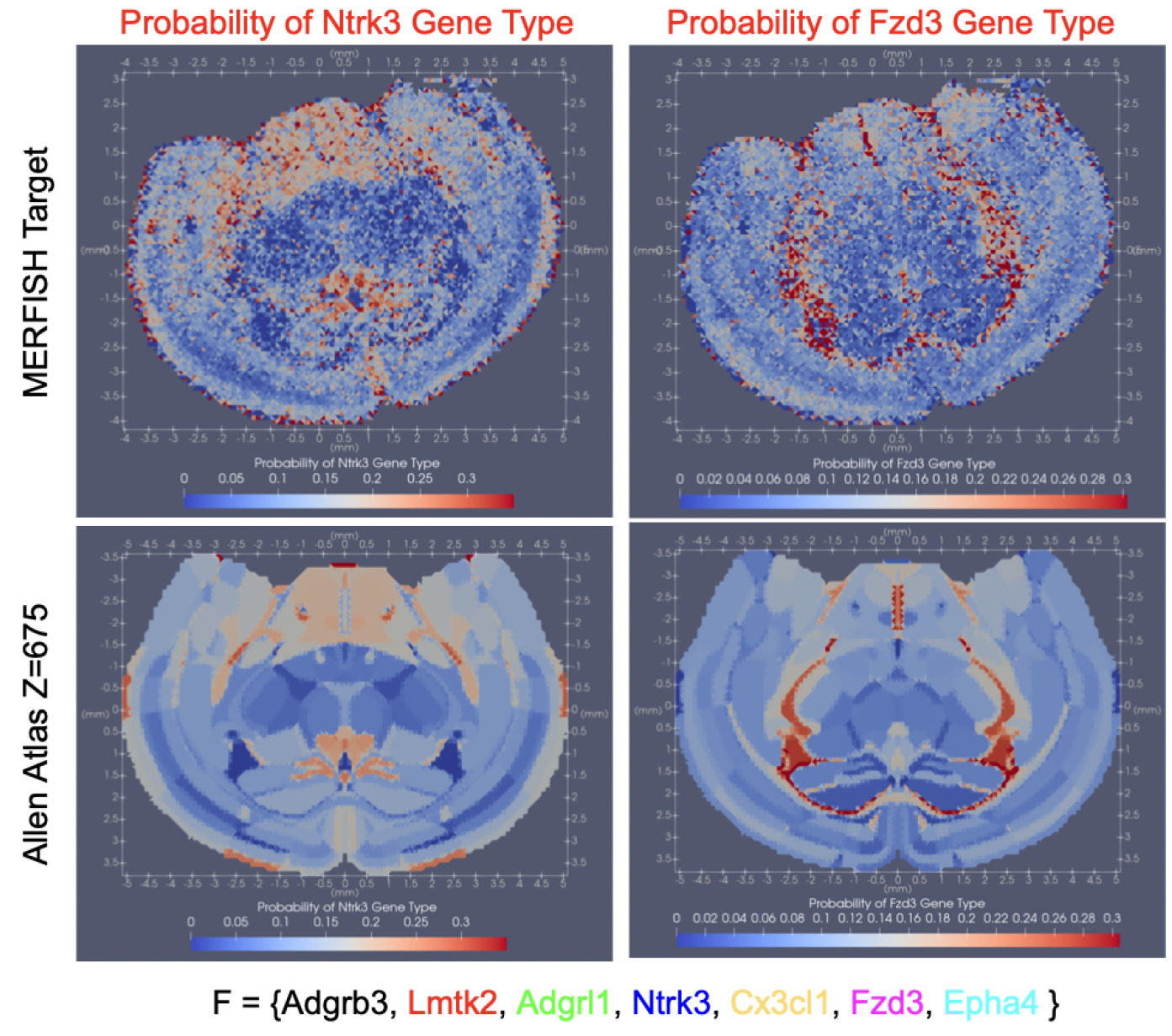
Gene type probabilities per cell for MERFISH section (top) for two genes *Ntrk3* gene type (left) and *Fzd3* gene type (right) out of a selected subset of 7 genes with high spatial discriminance according to mutual information score (see Section 4.4). Bottom shows estimated probabilities, ***ζ***^*π*^, for corresponding coronal Allen slice *Z* = 675.

### 2.5 Stability of Geometric Transformations Across Varying Feature Spaces

The previous section demonstrated the efficacy and flexibility of our algorithm at mapping cartoon atlases to a molecular target in the settings of that target carrying either gene-based or cell-based functional information. The assumed biological correlation between cell type and pattern of gene expression implies that signals of variation across cell types at the scale of microns should also exist across gene types at the scale of nanometers. Consequently, we might expect similar spatial deformation of a tissue scale atlas in mapping onto the same geometric target, but with conditional feature distributions defined over either gene or cell types, with partition boundaries deforming to match regions of homogeneity that would be roughly consistent across genes and cells.

Figure 6 shows the diffeomorphisms estimated for mapping Allen atlas sections at *Z* = 890 and *Z* = 675 onto two MERFISH target sections carrying three different feature spaces constructed from the same starting spatial transcriptomics data. Comparing left to middle and right columns, we see similar geometric transformations, *φ*, estimated to bring atlas onto target image varifold carrying cell type (left) versus gene type (middle, right) functional features. Regions of shrinkage (blue) versus expansion (red) occur in consistent areas across the different cases, and the magnitude of that change, as measured by the determinant jacobian, |*Dφ*|, is also similar in each case. Furthermore, we illustrate the effect of using two different subsets of 7 spatially discriminating gene types as the feature space. The first carries a high score based on Moran’s I (middle) and the second with a high mutual information score (right), as described in Sections 2.3 and 4.4. Here again, we observe similarity in the geometric mappings estimated for carrying atlas onto target between these two independent feature spaces. Hence, the manifest stability in the geometric mappings jointly estimated with the feature laws, 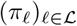, over three different feature spaces supports the stability of our alternating algorithm in the face of different numbers and types of features, but also speaks to the stability of the biological organization across tissue, cellular, and molecular scales.

**Fig. 6.**
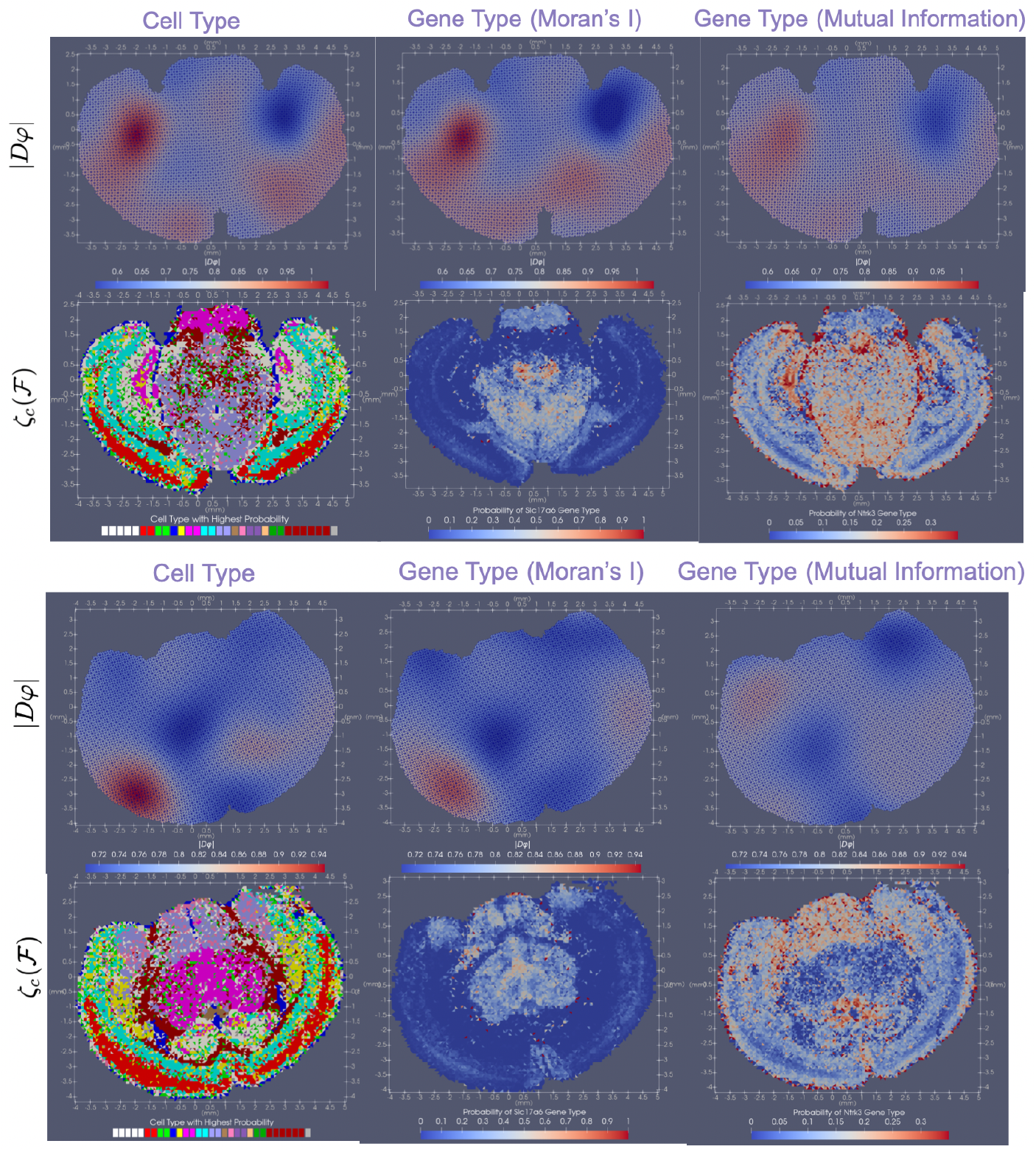
Diffeomorphism mappings of Allen atlas sections *Z* = 890 (top) and *Z* = 675 (bottom) to corresponding MERFISH sections for different feature spaces (e.g. cell types or gene types for a chosen subset of 7 genes). Top rows show the determinant of the Jacobian of the mapping |*Dφ*_1_| displaying areas of expansion (red) and contraction (blue). Bottom rows display different features on MERFISH sections including cell type (left), gene type in 7-gene subset selected with Moran’s I (middle), and gene type in 7-gene subset selected with Mutual Information (right). Cell types plotted as that with maximum probability; gene types plotted as probabilities for one specific gene in each subset.

### 2.6 Generalizing the Methodology to Compare Atlas to Atlas

The variational problem we solve via quadratic programming and LDDMM mapping between coordinates in (5) is universal in the sense that varifold representations can not only be used to represent the MERFISH sections of cellular and molecular data, as described in sections 2.3 and 2.4, but as well can be used to represent atlases, themselves. This allows us to map multiple atlas ontologies, one to another.

This is important because widespread variations in brain atlas ontologies have been developed to represent the molecular, chemical, genetic, and electrophysiological signals being measured across institutions. With different levels of granularity and different intended applications, multiple atlases per species now exist and are continuing to emerge [1, 2, 41, 42, 43, 44]. While some atlases have been defined in the same coordinate framework—often achieved through existing methods of image registration or manual alignment [28]— many exist in different coordinate frameworks. Together with mismatches in number, type, and positioning of partitions, this poses a challenge not only to the evaluation of each atlas ontology’s fit to a molecular target, but also the ready comparison of atlas to atlas and the establishment of a clear metric of similarity between them.

Figure 7 shows the results of mapping corresponding sections of both the ARA and Kim Lab Developmental atlas [43] to the cell-segmented MERFISH section of Figure 4. The images of predicted cell types with the highest probability (left column) for each compartment are shown for each ontology in the left column. The areas of the hippocampus (dashed circle) and striatum and amygdala (arrow) are partitioned with different levels of granularity. This leads to different optimal geometric transformations, as characterized by the determinant Jacobian (middle column), and different predicted cell type distributions (right column). Though both atlases are published as geometrically aligned [43], the diffeomorphism solving the variational problem transforms geometrically the homogenous regions between the atlas and target. Hence, regions of the amygdala and striatum undergo significant contraction in the optimal mapping of Kim but not Allen atlas to MERFISH given the partitioning of this region into fewer and thus larger presumed homogenous regions in the Kim atlas. The right column exhibits the entropy of the distributions over cell types estimated for each region in each atlas. Here, the hippocampus is more finely partitioned in the Kim atlas, which yields lower entropy distributions over cell types than in those estimated for the Allen atlas.

**Fig. 7.**
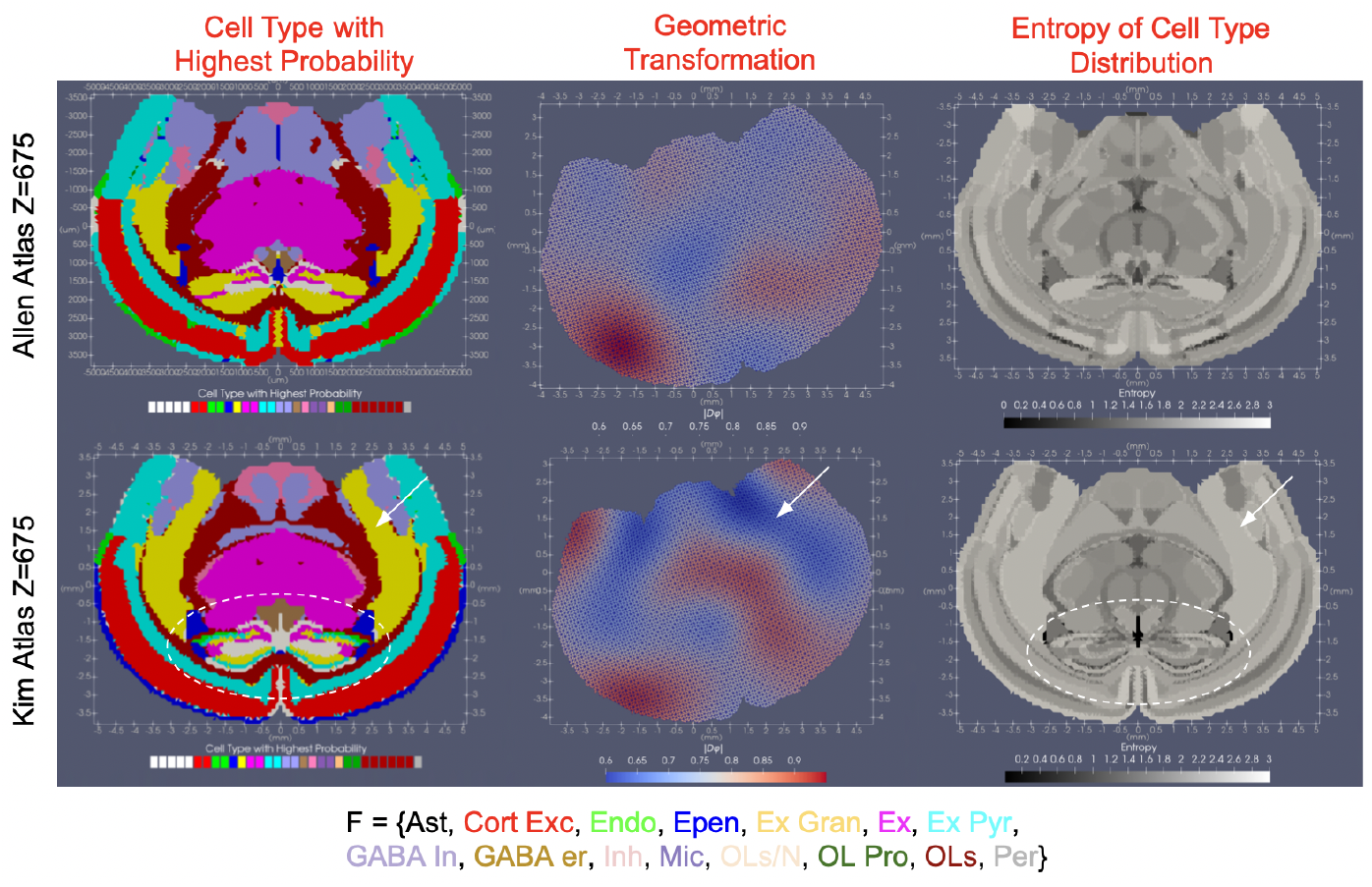
Comparison of mappings of *Z* section 675 of ARA (top) and Kim atlas (bottom) to cell-segmented MERFISH section. Left shows cell type with highest probability per simplex in each atlas. Middle shows estimated geometric transformation, *φ*_1_, in each setting applied to each atlas, with areas of expansion (red) and contraction (blue) as measured by the determinant Jacobian, |*Dφ*_1_|, of each mapping. Right shows entropy of estimated cell type distribution per simplex in atlas. Circled area of hippocampus and arrow pointing to area of amygdala and striatium highlight differences in estimated mappings for each atlas.

The universality of the variational problem allows for direct mapping across atlas ontologies. Here, atlases are taken to have constant density, *α*^min^ = *α*^max^ = 1. Thus, the mapping problem from atlas with ontology 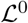 onto the target with ontology 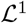 optimizes over the feature laws, 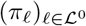, with the target atlas ontology 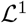 taken as the target feature space,

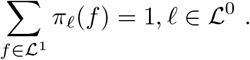

The joint estimation of geometric transformation, *φ* and conditional feature laws, 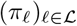 in our mapping methodology offers two modes of quantitative comparison of these atlas ontologies. First, as in classical image setting of LDDMM, the determinant jacobian, |*Dφ*|, of the estimated diffeomorphism, can be used as metric of how similar the atlas ontologies are, reflective of how much boundaries of partitions move to maximize overlap between homogenous regions. However, unlike in classical image settings, the estimation of the additional family of feature laws here affords a second metric of similarity with computation of the entropy of the estimated conditional feature distributions.

Figure 8 shows the results of mapping one mouse atlas ontology to another with the *Z* section 680 in the ARA mapped to the corresponding section in the Kim Lab Developmental atlas (top row) and vice versa (bottom row). The leftmost column depicts the geometry of the section under each ontology, with the Allen section hosting ≈ 140 independent regions and the Kim section ≈ 80. In this setting, both atlases are published in the same coordinate framework, giving *φ* = *Id* and thus, highlighting, instead, the estimated distributions over the other ontologies. The middle column depicts the estimated conditional probability distributions, ***ζ***^*π*^, for each atlas section over the other atlas section’s ontology. The label with the highest probability in these distributions is plotted for each simplex in the mesh and which is consistent across each partition of each original atlas, given the homogeneity assumption in our model (i.e. a single *π*_ℓ_ for each 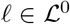). The comparatively larger set of labels in the Allen ontology results in labels being omitted from the corresponding estimated set of labels on the Kim ontology section (middle column, bottom row) while multiple regions in the Allen ontology carry the same most probable region in the Kim ontology. The right column of Figure 8 captures this difference in depicting the entropy of the estimated conditional feature distributions, 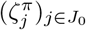, for each simplex of the mesh. The entropy of the distributions estimated for the Kim ontology over the Allen ontology (bottom) is on average, higher, than that of the distributions estimated for the Allen ontology (top), with probability mass distributed across ~ 5 – 7 different Allen regions for each Kim region of cortex. Nevertheless, we see close to 1:1 correspondence between Allen and Kim labels in the center section of the slice, where entropy of the estimated distributions is near 0.

**Fig. 8.**
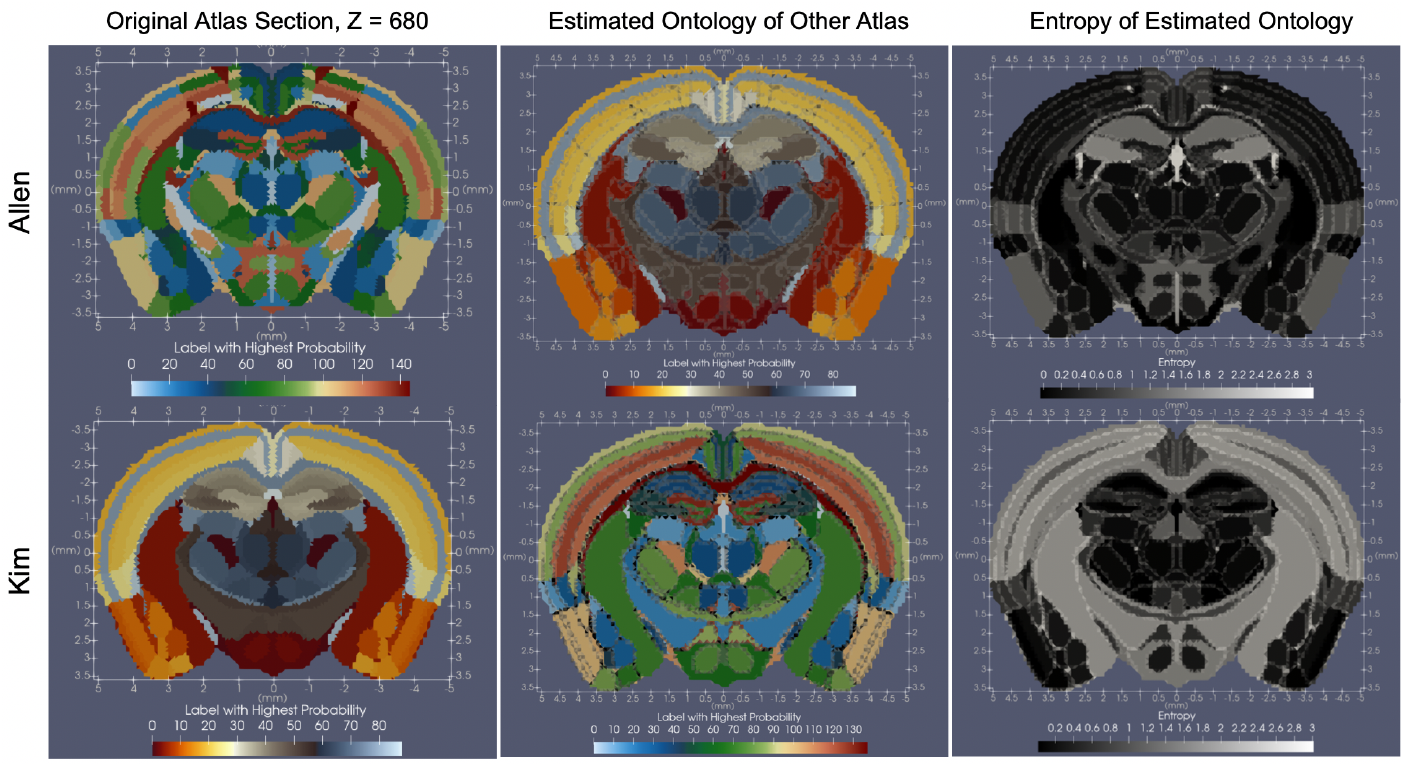
Original and predicted ontologies for Allen (top) and Kim (bottom) atlases. Left column illustrates original ontologies. Middle column illustrates Allen atlas geometry with Kim atlas ontology (top) and Kim atlas geometry with Allen atlas ontology (bottom). Right column shows entropy of predicted ontologies, with higher entropy values (light) indicating less 1:1 correspondence between ontologies.

Atlas ontologies can be mapped not just within species but also across them, where both geometric transformations and estimated ontology distributions, together reflect metrics of comparison between the two. Figure 9 shows the mapping of coronal section, *Z* = 537, in the ARA to a coronal section, *Z* = 628 in the Waxholm Rat Brain Atlas [44], with both sections chosen to correspond as sections through the anterior commissure. The left column shows both atlas ontologies with 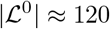 for the the Allen atlas section and 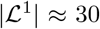 for the Waxholm atlas section. The middle column depicts the initial differences in size and shape (top) between the two tissue sections. After scaling the volume of the mouse brain by 1.5, additional deformation, with magnitude given by the determinant jacobian, |*Dφ*|, distorts both internal and external tissue boundaries to align homogeneous regions in each atlas, such as cingulate area to cingulate area (white arrow). Estimated distributions over the Waxholm ontology labels for each region in the Allen atlas are shown in the right column, summarized by the maximum probability label (bottom) and measures of entropy (top), which highlight in gray, Allen regions mapping to ≈ 3 – 4 Waxholm regions versus those in black achieving 1:1 correspondence.

**Fig. 9.**
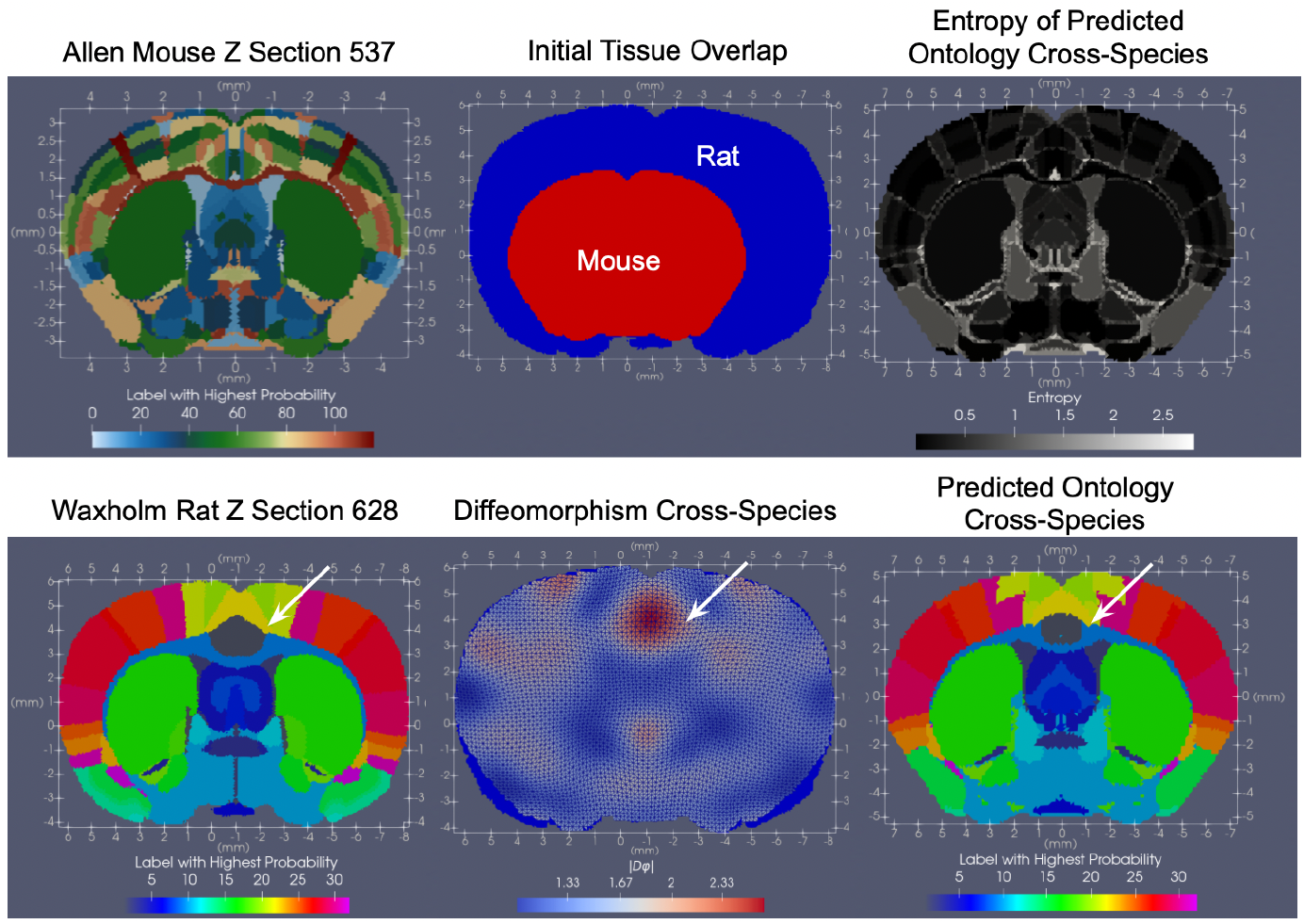
Results of mapping coronal section *Z* = 537 of ARA to corresponding coronal section of Waxholm Rat Brain Atlas at *Z* = 628, both chosen to be through the anterior commissure. Left shows both original atlas sections. Middle column shows initial tissue overlap between mouse and rat section (top) and resulting overlap following action of estimated diffeomorphism on mouse tissue (bottom). Determinant of Jacobian highlights areas of expansion (red) and contraction (blue) in mouse section deforming to match rat section, with white arrow highlighting expansion in cingulate area needed to match region in mouse to corresponding region in rat. Right column shows entropy for each mouse region’s predicted distribution of rat labels (top) and predicted rat label with highest probability (bottom).

## 3 Discussion

We have introduced, here, a universal method for mapping tissue scale, ‘cartoon’ atlases to molecular and cellular datasets arising in the context of emerging transcriptomics technologies. We root our method in the modeling of each object as a mesh-based image varifold, as previously described [27], and outline an alternating algorithm that simultaneously incorporates the classical deformation tools of LDDMM [29] with quadratic programming to jointly estimate an optimal geometric transformation and conditional feature law that maps atlas onto target.

As presented, our method fills a current need, as highlighted previously [35], for universal tools that can integrate the diverse types and large quantities of data emerging from the evolution of both transcriptomics and imaging technologies over the last decade. With each technology generating a slightly different perspective and different set of animal or human samples to compare, a method that can stably handle the format of past, current, and future datasets will be paramount to integrate both new findings with the vast number of datastores currently available across institutes. The image varifold framework used here is general enough to model emerging transcriptional data from both image-based and spot-resolution technologies and classical imaging data (as demonstrated in our atlas-to-atlas mappings). Therefore, it provides a gateway for comparing data historically curated through immunohistochemistry, MRI, and other techniques in addition to the emerging transcriptomics methods.

In parallel to the development and dispersion of diverse molecular datasets, there has been continued development on the side of reference atlases to reflect trends in these new measures and integrate these trends across even more samples of particular species. Our method offers a tool for re-examining and comparing existing atlas ontologies in the context of new data [35], and serves as a means for developing new atlases in the future. As described in Section 2.6, examination of the mappings achieved between different atlases and the same molecular target offers an indirect comparison between atlases in the context of a particular molecular setting. However, this comparison can also be made directly in a context-independent setting by harnessing our method to map atlas to atlas. In the field of evolutionary biology, for instance, our method could aid in the mapping and comparison of atlases across species [45, 46] and in the field of developmental neurobiology, the available atlases of the brain at different stages of development [42, 41]. With regard to atlas refinement and creation, the invertibility of the estimated diffeomorphism in the setting of mapping atlas to molecular target, enables the carrying of each target into the same coordinate space of the atlas. Here, the molecular and cellular scale raw reads could be averaged across individual samples, thus providing a mode for defining new atlas segmentation schemes of homogenous regions across these samples.

While the results presented here survey a wide variety of potential applications of our method to mapping atlas modalities to diverse targets, there remain uncertainties and potential modes of improvement that are the subject of current and future work. First, we have presented results mapping 2D sections of 3D atlases to corresponding 2D sections of MERFISH data. The Allen MERFISH data showcased here is part of an entire set of serial sections that span the whole brain. Consequently, we are optimizing our method to compute mappings of atlas to molecular target in 3D, where both added dimensions and added magnitudes of data contribute to the theoretical and computational complexity of the problem. Indeed, with ≈ 6 billion individual transcripts measured across the span of the brain, treatment of this data as a regular lattice image would require on the order of 1000 billion voxels at 1 *μ*m resolution, which is coarser than that needed even to resolve two molecules of mRNA. Hence, it becomes even more vital to treat such data in the particle setting, as presented here, where we capture the sparsity and irregularity of the data in modeling it effectively in its lowest dimension, as 6 billion individual particle measures. Second, though we have highlighted both gene-based and cell-based datasets achieved with image-based MERFISH technologies, we are currently investigating the use of our method to map data from spot-resolution technologies such as SlideSeq [47] and additional image-based technologies such as BarSeq [48], which introduce variations in both the number of genes measured and the scope of tissue (whole versus hemi-brain) measured.

Finally, we emphasize that central to the model posed here is the underlying assumption that each compartment has a homogeneous distribution over molecular features that is stationary with respect to space. This assumption holds in many settings, as we might expect, given the inherent construction of atlases often to delineate regions of particular cell types and thus, where we see a set of predominant cell types or gene types consistently across the region in the molecular scale data, as in Figures 4 and 5. However, we also see examples where this homogeneity assumption may not be appropriate. An example of this is seen in Figure 3 where the expression of *Trp53i11* appears to be distributed along a decreasing gradient medial to lateral within the striatum. Notably, the results presented here reflect a particular balance between expected deformation and this homogeneity assumption, imposed by the relative weighting of the separate terms in the cost function. Current work at controlling this balance further includes the addition of a term controlling the divergence of the vector field to the energy defined in the variational problem 5, which leads to solutions more robust to deformation within the interior of the tissue. Future work will also include more rigorous evaluation of how well this homogeneity assumption holds and the effect the given balance between the two terms might have in different settings.

## 4 Methods

### 4.1 Construction of Mesh-based Image Varifold for Different Modalities

As introduced in Section 2.2, we represent each image varifold object as a triangulated mesh. Each mesh is built from a collection of vertices, **x** = (*x_i_*)_*j*∈*I*_ with each *x_i_* ∈ ℝ^2^, here. Each simplex in the mesh is defined from the vertices denoted as *γ*(**x**) and is paired with a 3-tuple with components that index the vertices of the simplex, (*γ*(**x**), *c* = (*c*^1^, *c*^2^, *c*^3^) ∈ *I*^3^) and determine the center 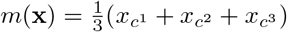. Each triangle simplex is defined by

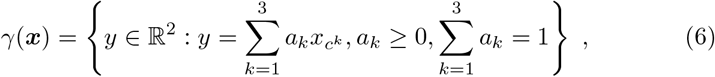

with positive orientation and volume 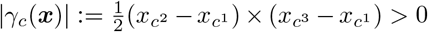.

The total mesh *τ* is the collection of vertices **x**, and simplices and centers 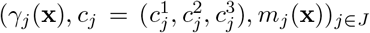 with the resolution determining the complexity as total numbers of vertices |*I*| and the number of simplices |*J*| in the mesh.

Meshes were constructed using Delauney triangulation [49] on a grid defined over the support of the starting dataset with the size of each square dictated by the input resolution. Varifold measures, ***α, ζ***, were associated to the simplices of the mesh following assignment of each individual data point (e.g. mRNA or cell read) into its single nearest simplex. Meshes were pruned of simplices that both contained fewer than 1 data point and existed outside the largest connected component of simplices containing at least one data point. In this manner, both for atlas images and transcriptomics data sets, resulting simplex meshes spanned the entire tissue foreground.

### 4.2 Molecular Scale Varifold Norm

To specify the image varifold norm for 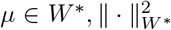, it suffices to provide the inner product between Diracs 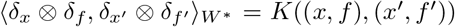, the right-hand side the kernel with for any weighted sum *μ* in Eqn. (1) then

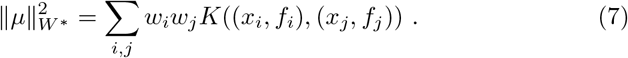

Throughout we use the kernel product *K*((*x, f*), (*x*′, *f*′)) = *K*_1_(*x, x*′)*K*_2_(*f, f*′) chosen as a Gaussian over physical space 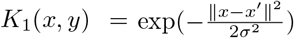 with *K*_2_(*f, f*′) = 1 if *f* = *f*′, 0 otherwise giving:

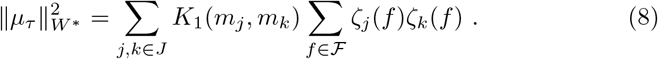

### 4.3 Alternating LDDMM and Quadratic Program Algorithm for Joint Optimization

For solving the variational problem of (5) we follow [27] using an alternating optimization, fixing the laws 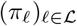 and optimizing over the control *v*(*t*), *t* ∈ [0, 1] and integrating it to generate the diffeomorphim *φ*_1_, then fixing the diffeomorphm and using quadratic programming to estimate the laws. The variational problem of (5) is optimized using LDDMM by flowing the atlas 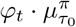 to minimize the target norm to the endpoint *μ_τ_*. Smoothness is enforced via the reproducing kernel Hilbert space norm on the control || · ||_*V*_ which controls the differentiability of the flow of vector fields, which is sufficient to guarantee an invertible diffeomorphic result [50]. Holding that fixed we alternately optimize (5) with respect to the laws 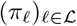 using quadratic programming, such as OSQP [51]. We loop until convergence.

**Algorithm 1.**
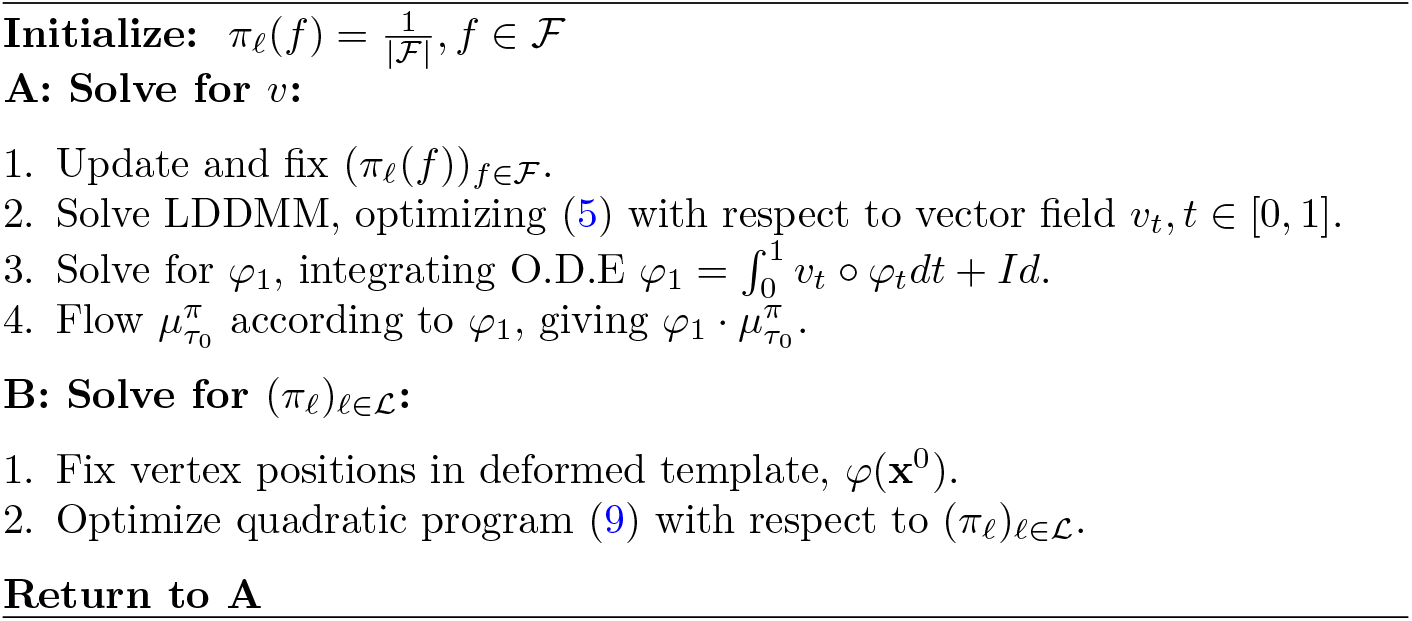

For the atlas, take the mesh *τ*_0_ with vertices **x**^0^ = (*x_i_*)_*i*∈*I*_0__ and with simplices and centers 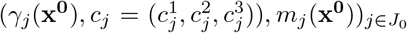. Estimated densities and conditional probabilities are denoted (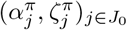. Define 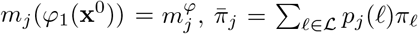, giving 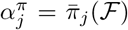. The quadratic program is given by:

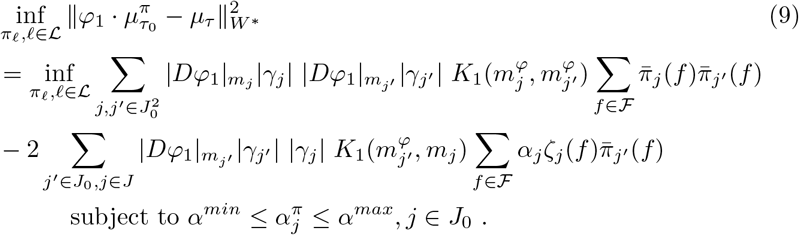

#### Remark 1

In the algorithm, we can use two approximations that are convenient. The first approximates the determinant of the Jacobian. Denoting 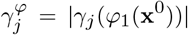, then approximating 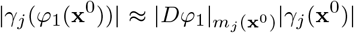 gives the simplified cost of the quadratic program:

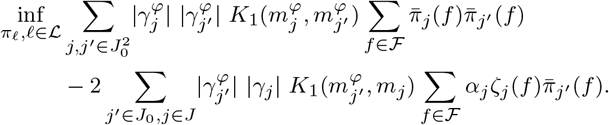

This can be simplified by representing the estimated laws 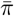 via the labels which have greatest area for the simplex. Defining the greedy maximizer map 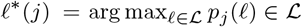, then the inner product can be approximated by

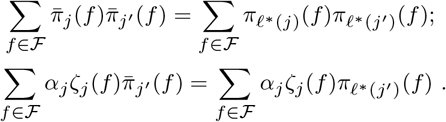

For simplex triangles within the interior of each atlas region, denoted *j* ∈ *J*_0_ \ *∂J*_0_, then 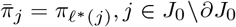 and these approximations are an equality for all interior pairs of vertices.

For all results shown, the template and target are initially aligned through separate estimation of rigid transformations (translation and rotation) and a single isotropic scaling applied to the template to bring the total area of the template to equal that of the target. Rigid transformations are estimated by minimizing the varifold normed difference Eqn. (9) between the rotated and translated template atlas 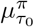 transformed to the target *μ_τ_*.

Everything being specified, gradient based optimization is performed until convergence or a specified number of iterations. In LDDMM, we use L-BFGS optimization combined with a line search using the Wolfe condition. In rigid registration, we directly optimize the varifold norm of the difference, also using the L-BFGS method.

### 4.4 Mutual Information Score for Discriminating Spatially Informative Genes

To deduce which genes are spatially variant with respect to their expression patterns, we assign to each gene a score based on mutual information. This score specifically measures the mutual information between a random variable, *M^g^*, that reflects the number of counts of gene *g* in a given neighborhood, and a random variable, *X*, that partitions this neighborhood vertically or horizontally into two domains. We describe, here, a method for computing this score particularly in settings of large amounts of data, where discretization is favorable for computational efficiency. This method, as illustrated in Figure 10, is applied for each gene independently on each measured section of tissue, where collective scores per gene be be garnered by tallying each gene’s score per section across the entire set of sections.

**Fig. 10.**
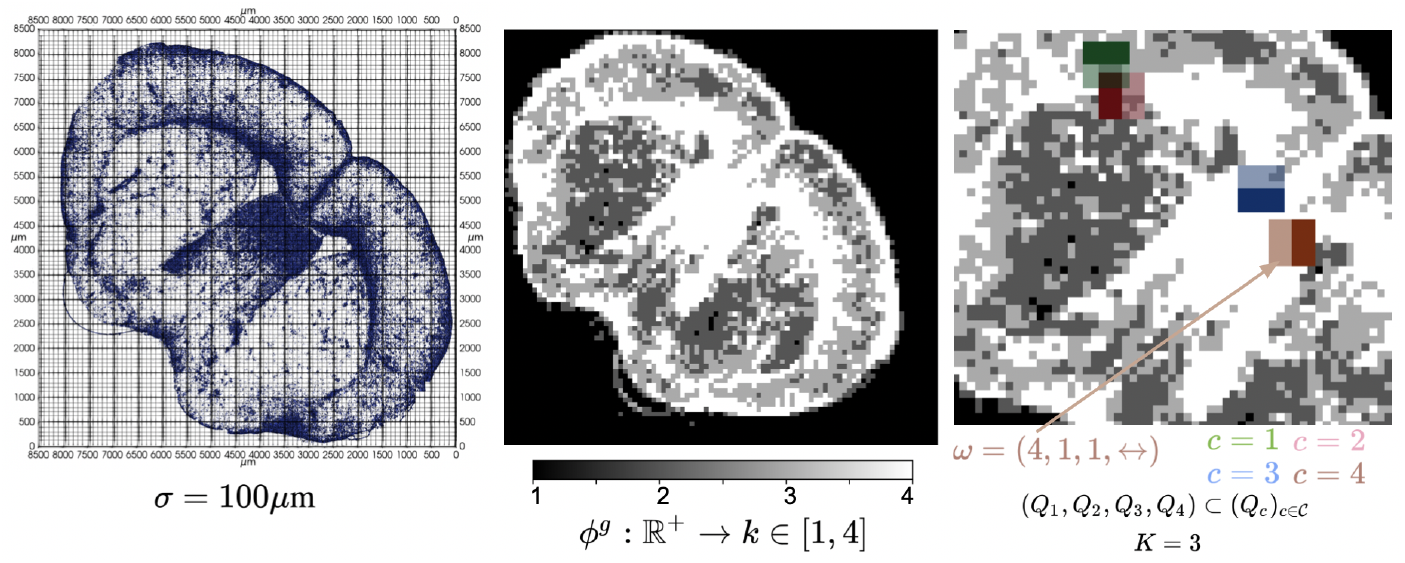
Steps in computing mutual information score for each gene in each tissue section. Left shows individual mRNA reads for gene *g* = Gfap. The support of the tissue is covered by a grid with squares of size *σ* × *σ*, with *σ* = 100*μ*m shown here. Middle shows output of binning function, *φ^g^* on the counts of gene *g* in each grid square, with *q* = 4. Right shows zoomed in portion of tissue with sample of 4 megacubes out of the entire set 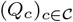. Example *ω* given for the individual grid square located at the bottom left corner of *Q*_4_.

The support of the tissue section is first covered by a grid, as shown in the left panel of Figure 10, with squares of size *σ* × *σ*. In the results shown in Sections 2.3 and 2.4, we choose *σ* = 50*μ*m. In each square, we compute the total number of mRNA expressed per each gene in that square, denoted by *N^g^* for gene *g.* Let *F^g^* (*t*) = *P*(*N^g^* ≤ *t*) be the cumulative distribution function for gene *g*, estimated from the empirical distribution of *N^g^* across all squares in our grid. We define the binning function 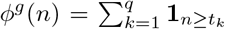 for *t_k_* = inf{*t* ≥ 0|*F^g^*(*t*) ≥ *k/q*} and with *k* ∈ [1,*q*] denoting the k-th q-quantile. This gives a discrete (normalized) value of mRNA counts for gene *g* in each square of the grid, as shown in the middle panel of Figure 10 for *g* = Gfap.

We define our discrete neighborhoods as megasquares, denoted 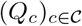, with each comprised of a continguous set of 2*K* × 2*K* grid squares. We consider all possible megasquares that can be defined across the grid, and index the squares within each megasquare by column index *i* = 1 · · · 2*K* and row index *j* = 1 ⋯; 2K, giving 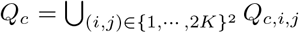. Finally, we define two partitioning schemes, denoted ↕ and ↔, corresponding to the partitioning of a megasquare into two equal vertical or two equal horizontal domains, each consequently containing 2*K*^2^ squares. The right panel of Figure 10 shows a sample of 4 megasquares from the entire set 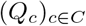 that cover the grid.

The random variables of interest, *X* and *M^g^* are specified as functions of *ω* = *(c,i,j,d*) ∈ Ω with 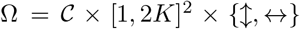, the set of all possible selections of megasquare, square within the megasquare, and partitioning of the megasquare. Specifically, we denote *C*(*ω*) = *c*, the index of the megasquare, *N^g^*(*ω*) the counts of gene *g* for the square *Q_c,i,j_* in megasquare, *c*, giving *M^g^*(*ω*) = *φ*(*N^g^*(*ω*)) ∈ [1,*q*], the q-quantile of the gene count, and *X*(*ω*) ∈ {*b,t,l,r*}, the partition *Q_c,i,j_* belongs to, dictated by direction *d* in *ω* as:

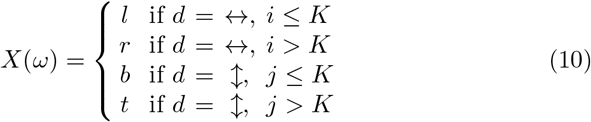

Choice of *ω* is made uniformly, with 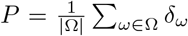. Our score is thus, the conditional mutual information between *X* and *M^g^* given *C*:

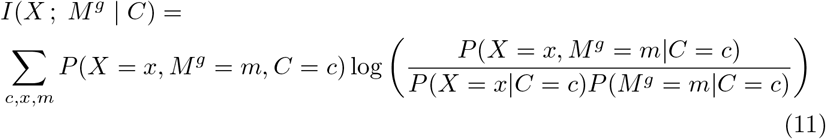

## Author Contributions

MM, AT, and LY developed the mathematical theory behind the manuscript. KS and MM drafted the manuscript. KS and LY generated codes for algorithms described in the manuscript. KS created all figures in the manuscript. MK, LN, and HZ generated serial MERFISH data. MA and JF annotated cell types for cell-segmented MERFISH data. YK created the reference atlas analyzed here with the Allen reference atlas. All authors contributed to the editing of the final manuscript.

## Materials and Correspondence

Questions, comments, and requests should be directed to corresponding authors, Kaitlin Stouffer and Michael Miller.

## Data Availability

Serial MERFISH sections from the Allen Institute were produced under the BRAIN Initiative Cell Census Network (BICCN, www.biccn.org, RRID:SCR_015820) and will be available at the Brain Image Library (BIL, https://www.brainimagelibrary.org/index.html) under doi https://doi.org/10.35077/g.610. Selected cell-segmented MERFISH sections were provided courtesy of Vizgen and together with cell type annotations are available upon request.

## Code Availability

Implementations of the algorithms described here can be found at: https://github.com/kstouff4/MeshLDDMMQP.

